# Correlation of Live Imaging and Fixed-Expansion Microscopy to study intracellular pathogens

**DOI:** 10.64898/2026.10.29.762944

**Authors:** Jennyfer Trouve, Aljoscha Janis Jung, Marek Basler

## Abstract

Expansion microscopy (ExM) enables high-resolution fluorescence imaging by physically enlarging biological samples, but its application to intracellular bacteria remains challenging. Here, we adapted the Ten-fold Robust Expansion Microscopy protocol to study host–pathogen interactions. Expansion of *Burkholderia thailandensis*-infected epithelial cells revealed previously unresolved membrane architectures associated with bacterial cell-to-cell dissemination. While host cells expanded approximately 9-fold, bacterial cells displayed incomplete and heterogeneous expansion. We found that fixation conditions strongly influence bacterial expansion, while presence of peptidoglycan partially limits expansion. To connect the membrane ultrastructures revealed by expansion with infection dynamics, we developed Live-to-Fixed Expansion microscopy (LiFEx), a workflow that correlates dynamic events observed by live-cell imaging with high-resolution cellular structures resolved by ExM. By correlating T6SS-5 assembly with subsequent high-resolution membrane architecture, LiFEx revealed membrane alterations developing within minutes following T6SS-5 assembly. Together, our approach enabled the investigation of dynamic host–pathogen interactions across complementary spatial and temporal scales.

## Introduction

Expansion microscopy (ExM) has emerged as a powerful and accessible approach to overcome the diffraction limit of conventional fluorescence microscopy. Rather than relying exclusively on specialized optical systems, ExM improves effective spatial resolution by physically enlarging biological specimens through anchoring of biomolecules to a swellable hydrogel^1^. Isotropic expansion requires homogenization of the sample, typically through enzymatic digestion or heat-induced denaturation, followed by swelling of the polymer network in deionized water. Since its initial development, numerous ExM variants have extended the approach to a broad range of biological specimens, including mammalian cells, tissues, plants, and microorganisms, with expansion factors ranging from approximately 4- to 20-fold^2–5^. Its compatibility with conventional fluorescence microscopes and diverse labeling strategies makes ExM particularly adapted to investigate cellular architecture at high spatial resolution.

Applying ExM to organisms surrounded by a protective cell wall nevertheless presents specific challenges, as this mechanically robust structure can restrict sample expansion. In all bacteria, the cell wall contains peptidoglycan (PG), a mesh-like structure composed of glycan strands cross-linked by short peptides that maintains mechanical integrity and protects the cell against osmotic pressure. Several bacterial ExM protocols therefore incorporate PG hydrolases, including lysozyme, mutanolysin, or species-specific enzymes, to facilitate sample homogenization and expansion^6–8^. However, PG composition, thickness, and architecture vary substantially between bacterial species^9^, resulting in different susceptibilities to enzymatic treatment and requiring species-specific protocol optimization^6^. In Gram-negative bacteria, the thin layer of PG is surrounded by an outer membrane (OM) that differs in lipid and protein composition from the inner membrane^10–12^. The OM is asymmetric, with a phospholipid-containing inner leaflet and a lipopolysaccharide (LPS)-rich outer leaflet, and is physically connected to the underlying PG.

To date, only a limited number of ExM protocols have been developed for bacterial imaging^6–8,13–18^, and only a subset have been tested on infected cells^6–8,13,14^. Most of these methods achieve approximately 4-fold expansion, while only a few protocols reach expansion factors of 8- to 11-fold^8,13,19^. To our knowledge, only one protocol has so far achieved a 10-fold expansion of intracellular bacteria^13^. However, this approach relies on a complex expansion strategy that limits its broad implementation in laboratories.

The final resolution achieved by super-resolution imaging is particularly important, as it directly determines the level of structural detail that can be resolved. This is particularly relevant for investigating host-cell membrane remodeling during the cell-to-cell spread of intracellular bacteria. Several intracellular bacteria, including *Listeria*, *Shigella*, and *Burkholderia*, exploit actin-based motility to move through the host-cell cytoplasm and reach the plasma membrane^20–22^. By pushing against the host membrane, these bacteria generate membrane protrusions that extend into neighboring cells. These protrusions can subsequently resolve into double-membrane vacuoles within recipient cells, from which bacteria escape using different species-specific strategies^20,23^. *Burkholderia thailandensis*, a surrogate for the highly pathogenic *B. pseudomallei*, employs a distinct mechanism involving its Type VI Secretion System 5 (T6SS-5)^24,25^, which is required for escape from these double-membrane compartments^26^. *Burkholderia* can also fire T6SS-5 while within membrane protrusions, leading to fusion of the plasma membranes of adjacent host cells. Repeated cell-cell fusion results in the formation of multinucleated giant cells (MNGCs)^24,27–29^. Although live-cell fluorescence microscopy has provided important insights into the dynamics of this dissemination process^26^, the membrane organization associated with these events remains difficult to resolve with conventional optical microscopy techniques.

Here, we optimized the Ten-fold Robust Expansion Microscopy (TREx) protocol^30,31^ to image mammalian cells infected with intracellular bacteria. We chose this protocol because (i) it achieves 10-fold expansion in a single expansion step with basic laboratory equipment, (ii) it is compatible with pan-protein and dense membrane labeling using commercial dies. Our approach enabled dense membrane and pan-protein labeling of infected cells and revealed membrane remodeling associated with *B. thailandensis* dissemination. However, we observed that bacteria expand substantially less than the surrounding host cells. We therefore investigated several parameters potentially limiting bacterial expansion, including PG degradation, outer membrane (OM) permeabilization, proteinase K digestion, and chemical fixation. OM destabilization and extensive proteolysis did not substantially reduce the heterogeneity of bacterial expansion. Surprisingly, PG degradation appeared to have minor impact on bacterial expansion however it increased expansion isotropy. In contrast, chemical fixation conditions strongly influenced bacterial expansion. Finally, we developed a correlative workflow that enables the same region of interest to be imaged with live-cell microscopy and subsequently with ExM. This approach, named LiFEx (Live-to-Fixed Expansion microscopy), bridges the gap between the high-resolution structural information provided by ExM and the dynamic biological context preceding fixation. Using LiFEx to correlate *B. thailandensis* T6SS-5 assembly with high-resolution membrane architecture revealed the delay between T6SS-5 assembly and the membrane fragmentation and protrusion lysis.

## Results

### Peptidoglycan limits proteinase K access to the bacterial cytoplasm

As there is no standardized approach for PG degradation in bacterial expansion microscopy, we first investigated whether the order of PG degradation and enzymatic sample denaturation impacted the degradation of bacterial cells. *B. thailandensis* expressing the fluorescent reporter mNeonGreen was grown *in vitro* and incubated for 2 h during exponential growth with HADA, a fluorescent D-amino acid that incorporates into bacterial PG^32,33^. Cells were subsequently washed and fixed with 4% paraformaldehyde (PFA) and 0.1% glutaraldehyde (GA). Fixed samples were then subjected to two successive enzymatic treatments: PG hydrolases and proteinase K, or the reverse order. PG degradation was carried out using a mixture of PG hydrolases (160 U/mL mutanolysin and 1 mg/mL lysozyme) for 3 h at 37°C in a magnesium-containing buffer. Protein digestion was performed with proteinase K (9 U/mL) for 3 h at 37°C. Finally, membranes were labeled overnight with mCLING^34^ at room temperature, and the samples were imaged on agarose pads (Figure 1).

**Figure 1:**
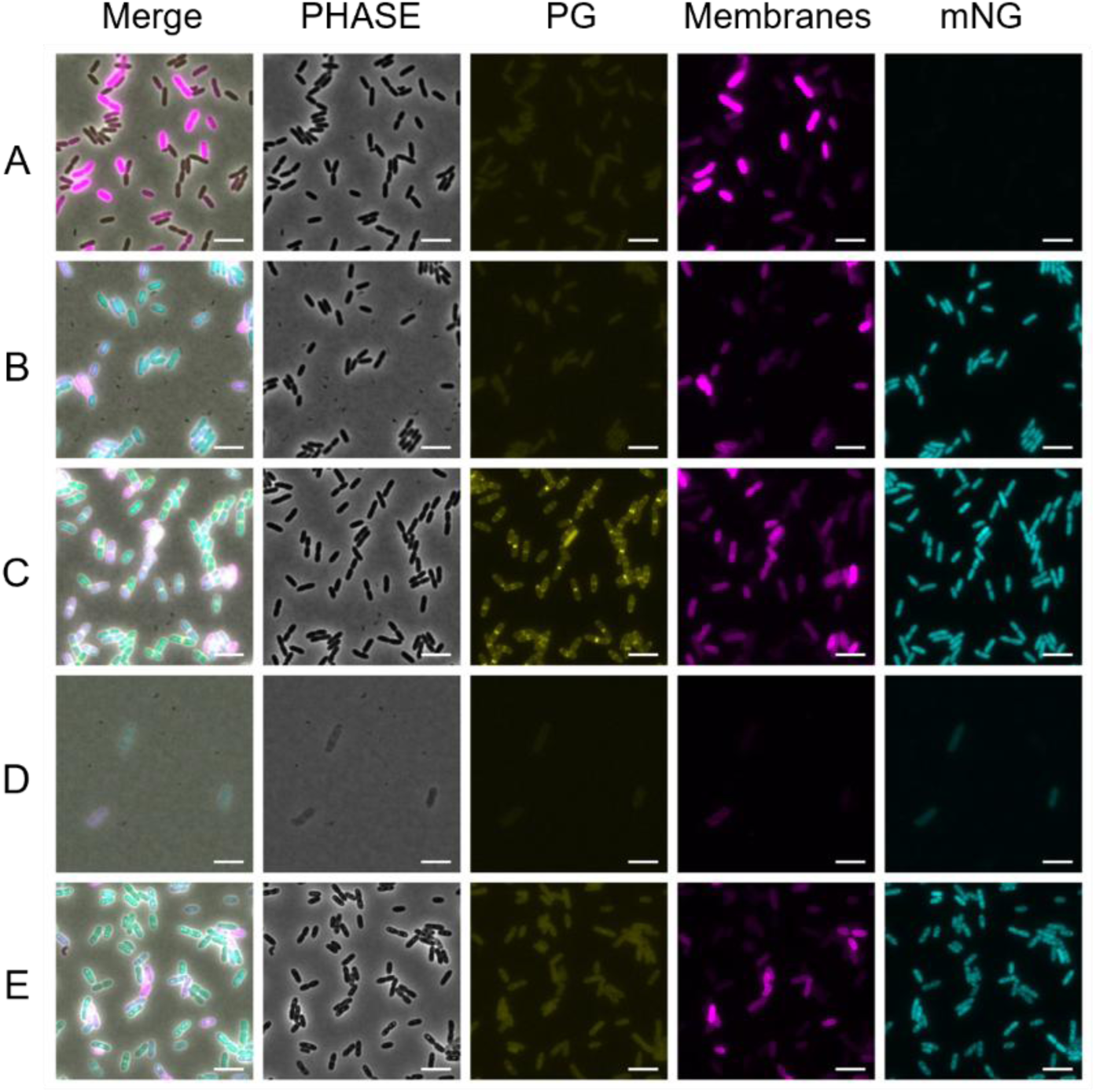
Peptidoglycan degradation facilitates proteinase K digestion of bacterial cells. Exponentially growing *B. thailandensis* were labeled with HADA, fixed, and subjected to successive enzymatic treatments. PG was degraded using mutanolysin and lysozyme, while proteins were digested using proteinase K. (A) *B. thailandensis* non-expressing mNeonGreen and unlabeled with HADA. (B) *B. thailandensis* expressing mNeonGreen and unlabeled with HADA. (C) *B. thailandensis* expressing mNeonGreen and labeled with HADA. (D) *B. thailandensis* expressing mNeonGreen, labeled with HADA and treated with PG hydrolases followed with proteinase K. (E) *B. thailandensis* expressing mNeonGreen, labeled with HADA and treated with proteinase K followed with PG hydrolases. Membranes are shown in magenta (mCLING), PG in yellow (HADA), and mNeonGreen in cyan. The phase-contrast images (PHASE) and the overlay of all channels (Merge) are also shown. Scale bar, 10 µm.

PG was completely degraded when PG hydrolases were applied prior to proteinase K, whereas substantial HADA-labeled PG signal remained when proteinase K was applied first (Figure 1 D-E). The order of the enzymatic treatments had a pronounced effect on protein degradation, particularly within the bacterial cytoplasm. Starting with PG degradation, resulted in reduced mNeonGreen signal upon proteinase K digestion (Figure 1D), suggesting that PG removal facilitates proteinase K access to the bacterial cytoplasm.

### ExM enables dense membrane and protein labeling of expanded infected cells

To visualize membrane ultrastructure during *B. thailandensis* infection of A549 cells, we augmented the existing TREx protocol with a PG degradation step. This protocol was applied to infected A549 cells as follows. We seeded adherent A549 cells on a round glass coverslip of 13 mm diameter, for 24 hours. We then infected the cells with *B. thailandensis* at a multiplicity of infection (MOI) of 100, for 17 hours prior to chemical fixation. Fixed samples were incubated overnight with fluorescent mCLING probes to stain the membranes. After permeabilization, the PG was degraded with a mix of mutanolysin and lysozyme in a magnesium-containing buffer, and the sample was further treated with the anchoring molecule AcX overnight. We placed the sample facing down on a monomer solution, which polymerized into a gel after incubation at 37°C. The sample inside the gel was then denatured (or homogenized), by enzymatic treatment using proteinase K. Finally, the DNA and proteins were stained with Hoechst 33342 and fluorescent NHS ester (“NHS” hereafter), respectively. To reach a complete expansion, the gel was placed in ddH2O in 3 consecutive baths for a total time of 24 hours.

With this procedure, we obtained expanded infected cells with a dense and continuous labeling of the host cytoplasmic membranes, a weaker membrane labeling for the host nucleus and the bacterial membranes (Figure 2B). The protein staining was consistent across the host cells. Interestingly, the NHS staining is higher in the bacteria than the host cytoplasm, potentially due to higher protein density in the bacterial cytoplasm. DNA was stained in both host nuclei and bacteria.

**Figure 2:**
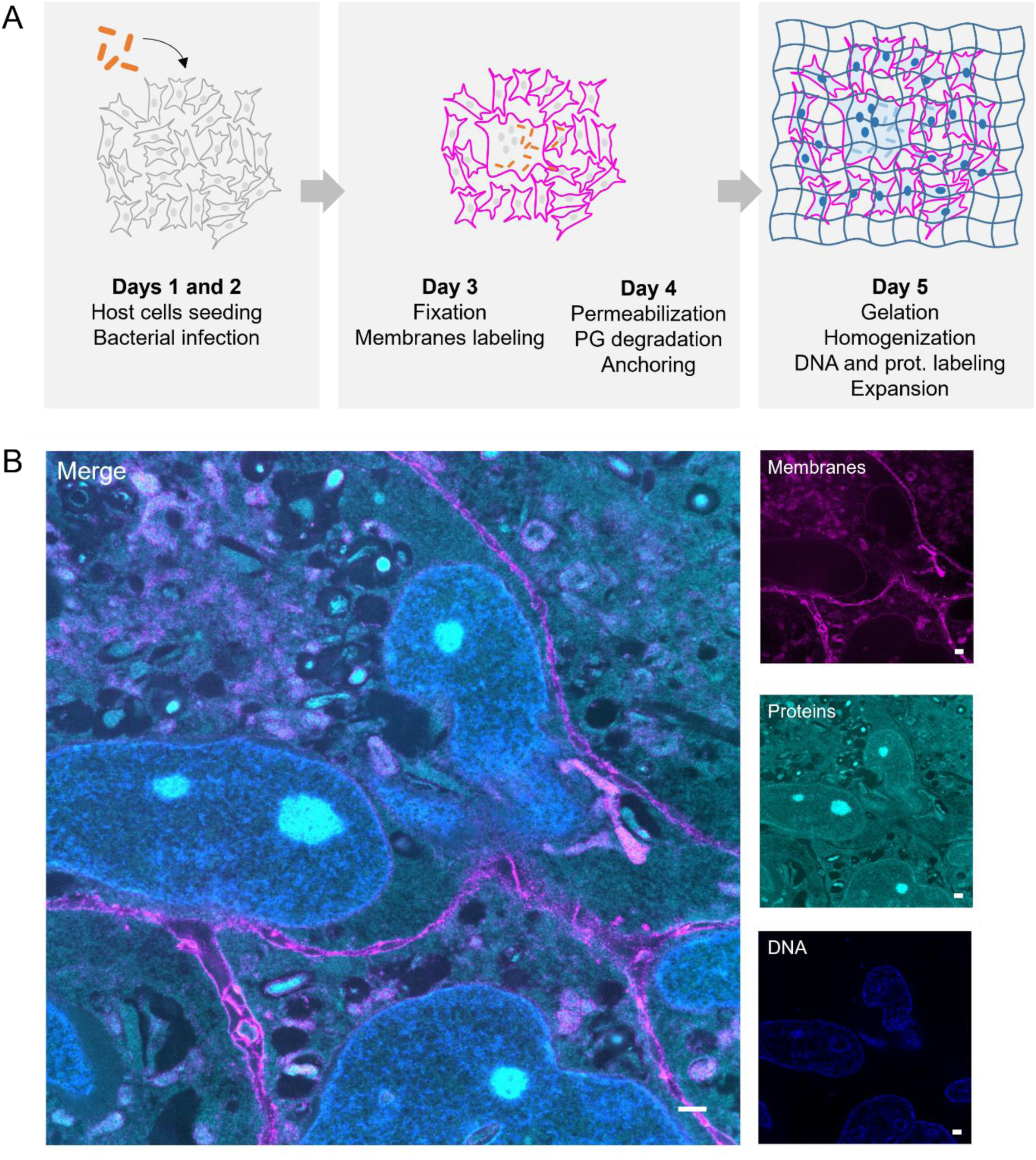
Optimized expansion workflow for analysis of host-pathogen interaction. **(A)** Schematic representation of the main steps of the protocol. **(B)** Overview of A549 cells infected with *B. thailandensis* (MOI 100, for 17h) after expansion. Membranes (mCLING) are shown in magenta, proteins (NHS) in cyan and DNA (Hoechst) in blue. Scale bar, 10 µm (post-ExM).

Expansion allows visualization of membrane lysis by *B. thailandensis*.

At later timepoints, from 17 hours post infection (hpi), intracellular *B. thailandensis* spreads between host cells by pushing the cytoplasmic membrane of the infected host cell, toward an adjacent cell. This actin-tail motility dependent process results in the formation of a double membrane protrusion^20–22^. The T6SS-5 of *B. thailandensis* is assembled almost exclusively inside these protrusions, resulting in the fusion of the membranes of the neighboring cells by a poorly understood mechanism. This repeated host membrane fusion leads to the formation of MNGCs, characteristic of *B. thailandensis* infection^24,27–29^. The protrusion formed by *B. thailandensis* can be also pinched by Dynamin-2 from the cytoplasmic membranes, forming a double membrane vacuole, from which the bacterium escapes in a T6SS-5-dependent manner^26^. Since MNGCs are easily recognized after expansion, we focused on imaging the periphery of these cells, where most membrane protrusions appear. Importantly, our expansion protocol allowed us to distinguish the two membranes forming the protrusions (Figure 3). We were also able to image different steps of the membrane protrusion formation and lysis. Some bacteria were surrounded by a protrusion that shows uniform labeling (Figure 3A), while others showed minor membrane staining discontinuities (Figure 3B). However, some bacteria were surrounded by protrusions with sections apparently lacking membrane signal (Figure 3C-D), and several other bacteria were rather surrounded by structures that resembled membrane fragments (Figure 3D). In addition, we also observed bacteria surrounded by double membrane vacuoles of different membrane staining patterns: some double membranes were regularly aligned (Figure 3E), while others formed irregular bulging (Figure 3F). Altogether, our expansion pipeline allowed high-resolution imaging of bacterial spread and membrane fragmentation caused by *Burkholderia*.

**Figure 3:**
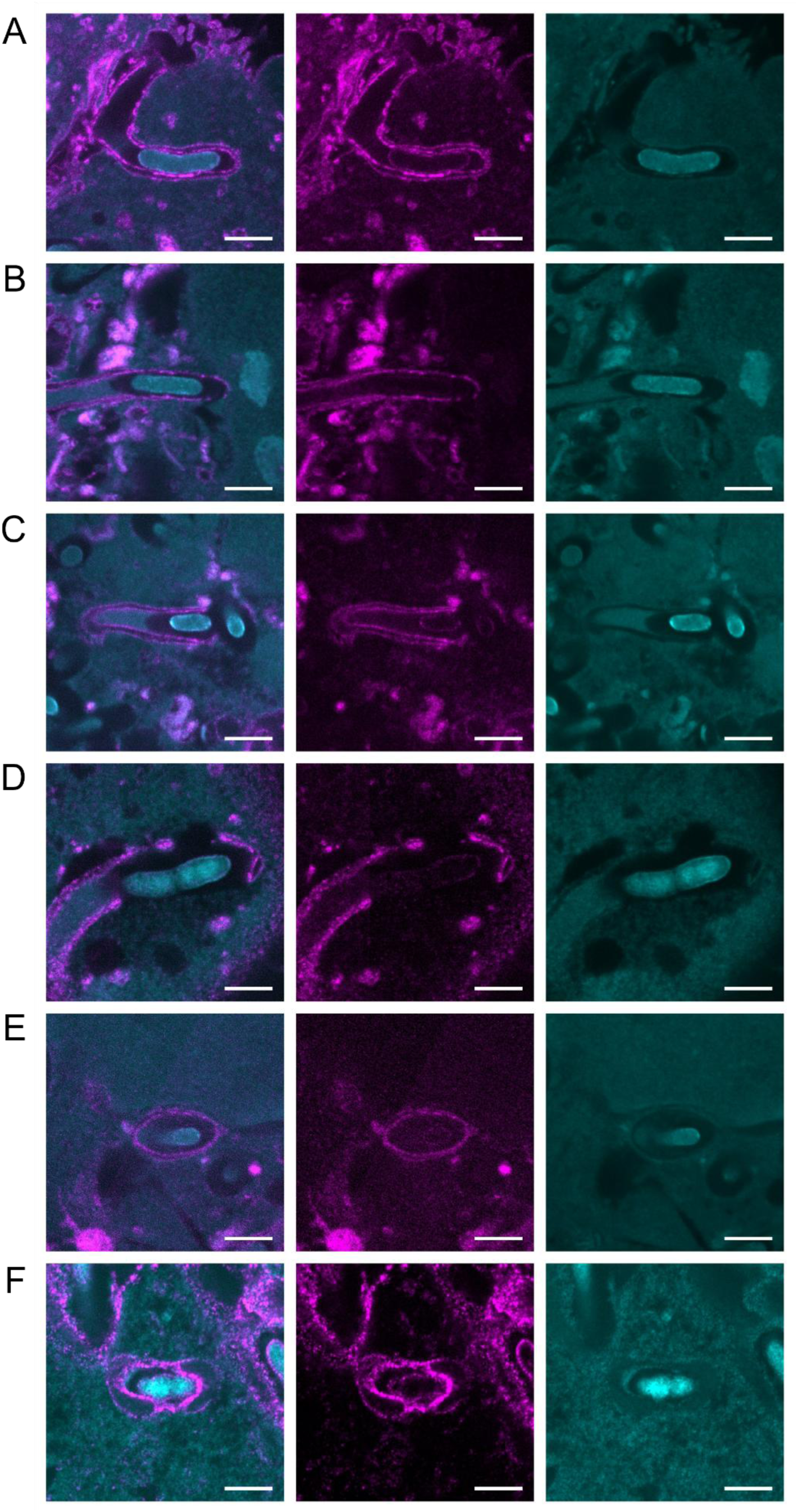
Expansion shows membrane protrusion utrastructures. **(A-F)** A549 cells were infected with *B. thailandensis* (MOI 100) for 17 h. After fixation, our optimized expansion protocol was applied, and the expanded samples were imaged using spinning disk confocal microscope. Membranes (mCLING) are shown in magenta and proteins (NHS) in cyan. Scale bar, 10 µm (post-ExM).

### Expansion of infected cells leads to heterogeneous bacterial expansion

When applying our expansion protocol to infected cells, we observed substantial variability in the expansion of individual bacteria within the same host cell. Poorly expanded bacteria were characterized by the absence of NHS signal surrounding the bacterial cell and increased NHS signal within the bacterial cytoplasm (Figure 3 and S1). To assess the extent of this heterogeneity, we compared the expansion factors of host cells and intracellular bacteria. Host-cell expansion was estimated by comparing the nuclear area before and after expansion using Hoechst staining. Bacterial expansion was determined by measuring the bacterial diameter, excluding division sites, before and after expansion based on membrane labeling. Using these measurements, we estimated an expansion factor of approximately 9× for host cells, and 5.5× for the intracellular bacteria. While the 9× expansion factor measured for host cells was consistent with the overall gel expansion, which typically ranged from 8- to 10-fold, the lower expansion factor measured for intracellular bacteria revealed a local expansion anisotropy associated with bacterial cells.

### Peptidoglycan influences but does not prevent bacterial expansion

To understand the incomplete expansion of intracellular bacteria, we first investigated whether the bacterial cell wall could represent a physical barrier to expansion. To determine the contribution of PG to bacterial expansion, we expanded infected cells with or without the PG degradation step (Figures 2A and 4A) and compared the post-expansion NHS labeling. Surprisingly, bacteria were able to expand in the absence of PG degradation, indicating that removal of PG is not strictly required for bacterial expansion. However, for some bacteria, the absence of PG degradation increased the expansion anisotropy along the transverse axis (width), resulting in elongated and thin bacteria (Figure 4A, white arrows). This indicates that PG can constrain bacterial expansion, particularly along the transverse axis, and that its degradation increases isotropic expansion but is not sufficient for achieving complete and homogeneous expansion of intracellular *B. thailandensis*.

**Figure 4:**
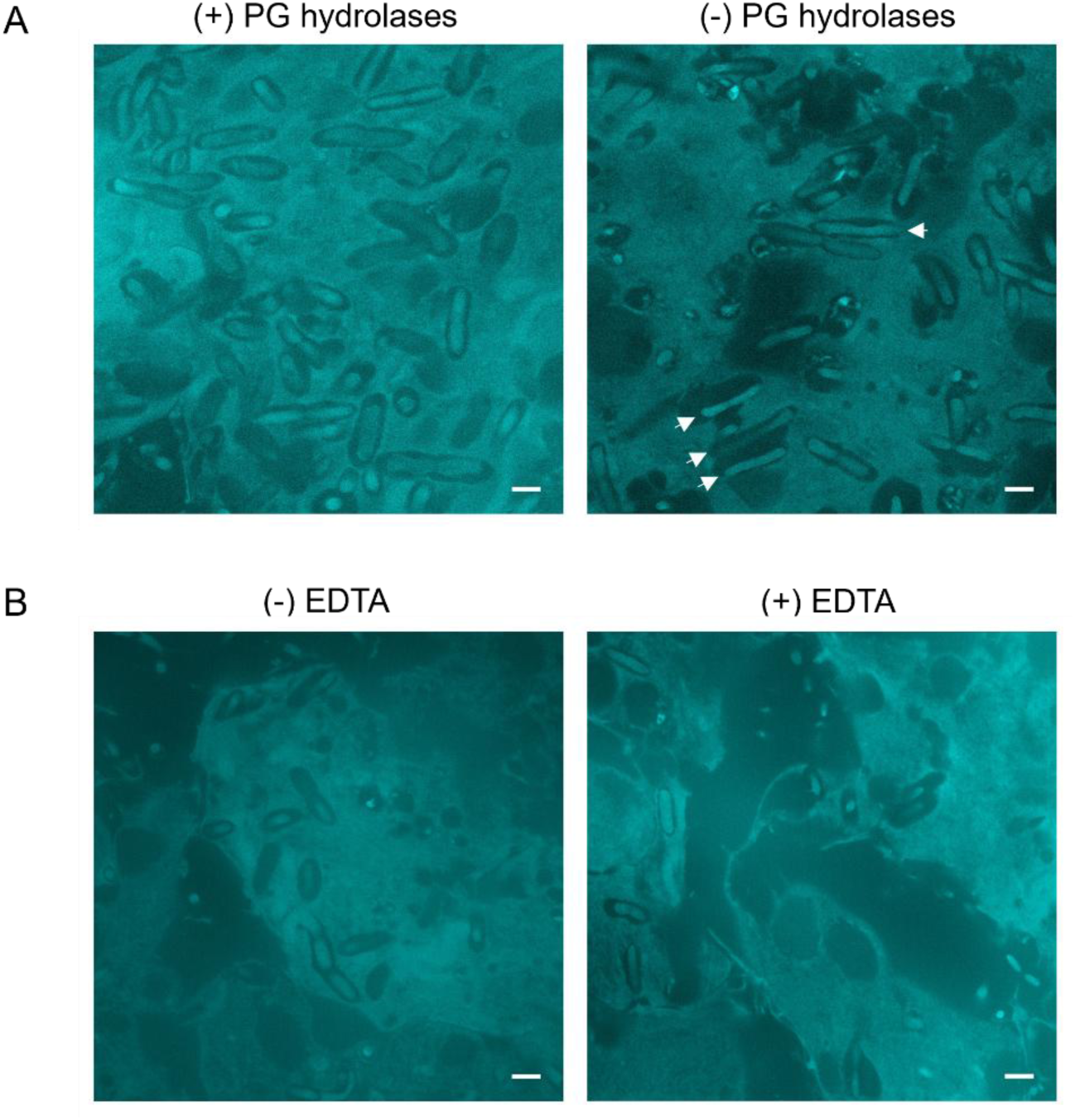
Effect of bacterial cell wall degradation on bacterial expansion. Expansion protocol was applied to A549 cells infected with *B. thailandensis* for 17 h with modification **(A)** in the PG degradation step or **(B)** by adding bacterial permeabilization step. **(A)** Comparison of expansion following PG degradation (left) and in absence of PG degradation step (right; white arrows indicate thin elongated bacteria). **(B)** Comparison of expansion without addition of EDTA (left) and addition of 5 mM EDTA prior to PG degradation (right). Protein (NHS) staining is shown in cyan. Scale bar, 10 µm (post-ExM).

We next investigated whether the OM could limit bacterial expansion. Divalent cations contribute to OM stability by promoting interactions between neighboring LPS molecules while chelation of these cations with EDTA destabilizes the OM and increases its permeability^35,36^. We therefore incubated fixed and permeabilized infected cells with 5 mM EDTA before proceeding with the remaining steps of the expansion protocol as described above (Figure 2A). No major differences in NHS labeling patterns were observed following EDTA treatment. Heterogeneous bacterial expansion persisted despite OM permeabilization, indicating that destabilization of the OM is insufficient to achieve uniform bacterial expansion to the same extent as host cells.

Together, these results indicate that the components of the bacterial envelope have a limited influence on bacterial expansion.

### Extended proteolysis fails to improve bacterial expansion

To determine whether the limited bacterial expansion resulted from incomplete protein digestion, we enhanced the proteolysis step by increasing either the concentration of proteinase K by threefold (from 9 to 27 U/mL) and the duration of the treatment (from 3h to 6h). Comparison of NHS staining across the different conditions revealed no noticeable improvement in bacterial expansion (Figure S2), indicating that bacterial expansion homogeneity cannot be improved by extended proteolysis.

### Bacterial expansion is sensitive to fixation conditions

To investigate the influence of chemical fixation on bacterial expansion, we tested different concentrations and combinations of PFA and GA before expanding infected A549 cells. Post-expansion NHS staining revealed pronounced differences in bacterial expansion patterns depending on the fixation conditions (Figure 5). However, we did not observe a clear relationship between fixative concentration and bacterial expansion: reducing the concentration of fixative did not consistently result in improved expansion. Instead, fixation with combinations of PFA and GA resulted in more homogeneous bacterial expansion than fixation with either PFA or GA alone. These observations indicate that bacterial expansion is highly sensitive to fixation conditions, however, this effect cannot be explained simply by the overall concentration of fixative.

**Figure 5:**
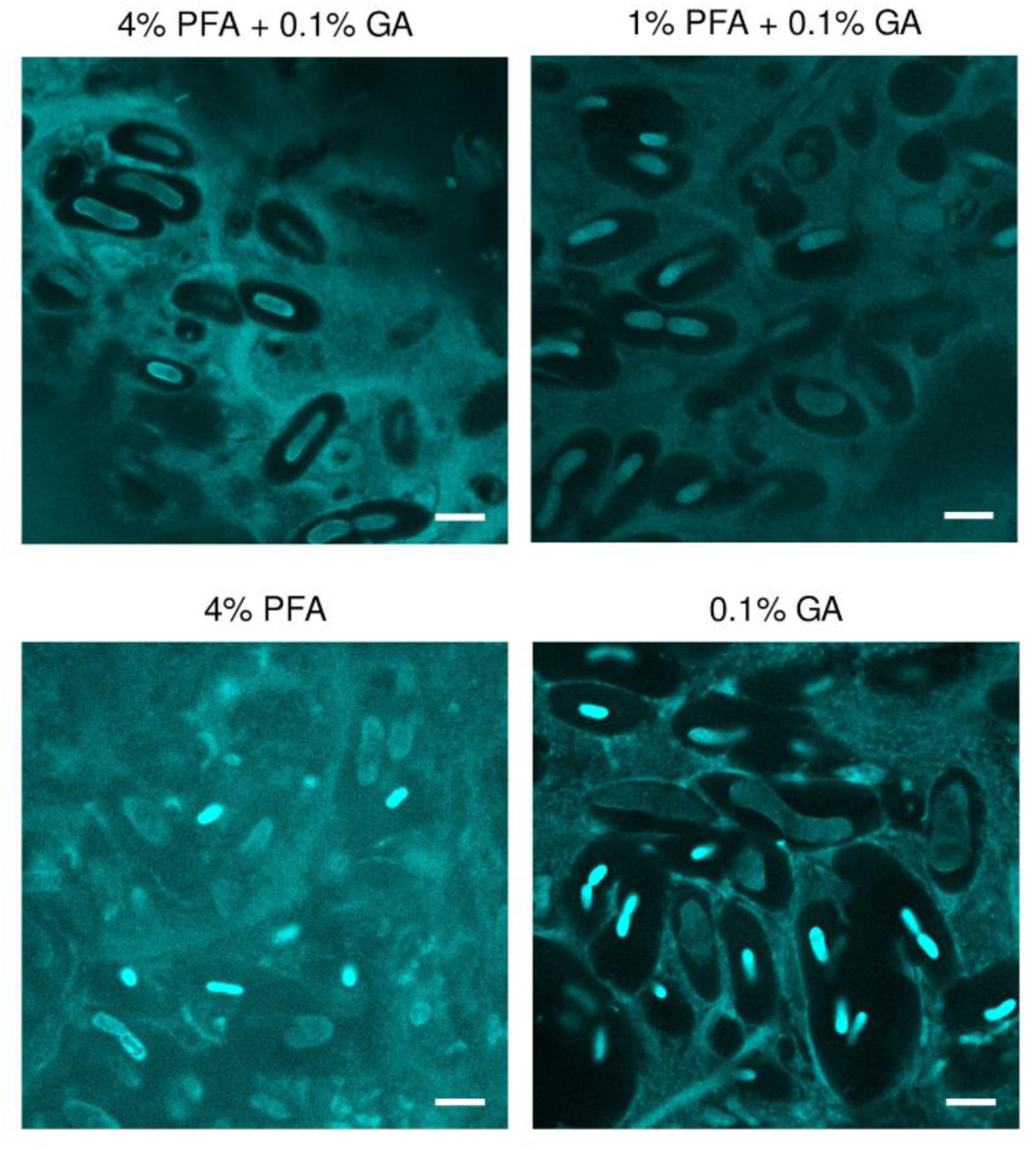
Fixation conditions influence bacterial expansion. A549 cells infected with *B. thailandensis* for 17 h were fixed using different concentrations and combinations of chemical fixatives, as indicated for each panel, before expansion. Protein staining (NHS) is shown in cyan. Scale bar, 10 µm (post-ExM).

### Correlation of live-cell dynamics with post-expansion cellular ultrastructure

ExM, like most super-resolution microscopy approaches, relies on fixed samples and therefore captures the structures present at the time of fixation, while cellular dynamics preceding fixation is not available. To bridge this gap, we developed a pipeline to correlate cellular dynamics observed by live-cell microscopy with the corresponding cellular ultrastructure subsequently resolved after expansion. We termed this approach LiFEx, for Live-to-Fixed Expansion microscopy (Figure 6A). Adherent A549 cells were seeded in 8-well glass-bottom µ-Slides for 24 h and subsequently infected for 17 h with *B. thailandensis* expressing the T6SS-5 sheath protein TssB-5 fused to mScarlet-I, at a MOI of 50. At 17 hpi, live cells were labeled with CellMask and Hoechst to visualize membranes and DNA, respectively. To facilitate matching expanded images to live-cell images, the observation was restricted to a defined 3×3 mm area marked on the bottom of the well. A region of interest (ROI), here corresponding to the interface between infected and uninfected cells where membrane protrusions form, was imaged by spinning-disk confocal microscopy using a 60× objective. Time-lapse images were acquired every 30 s for approximately 10–30 min.

**Figure 6:**
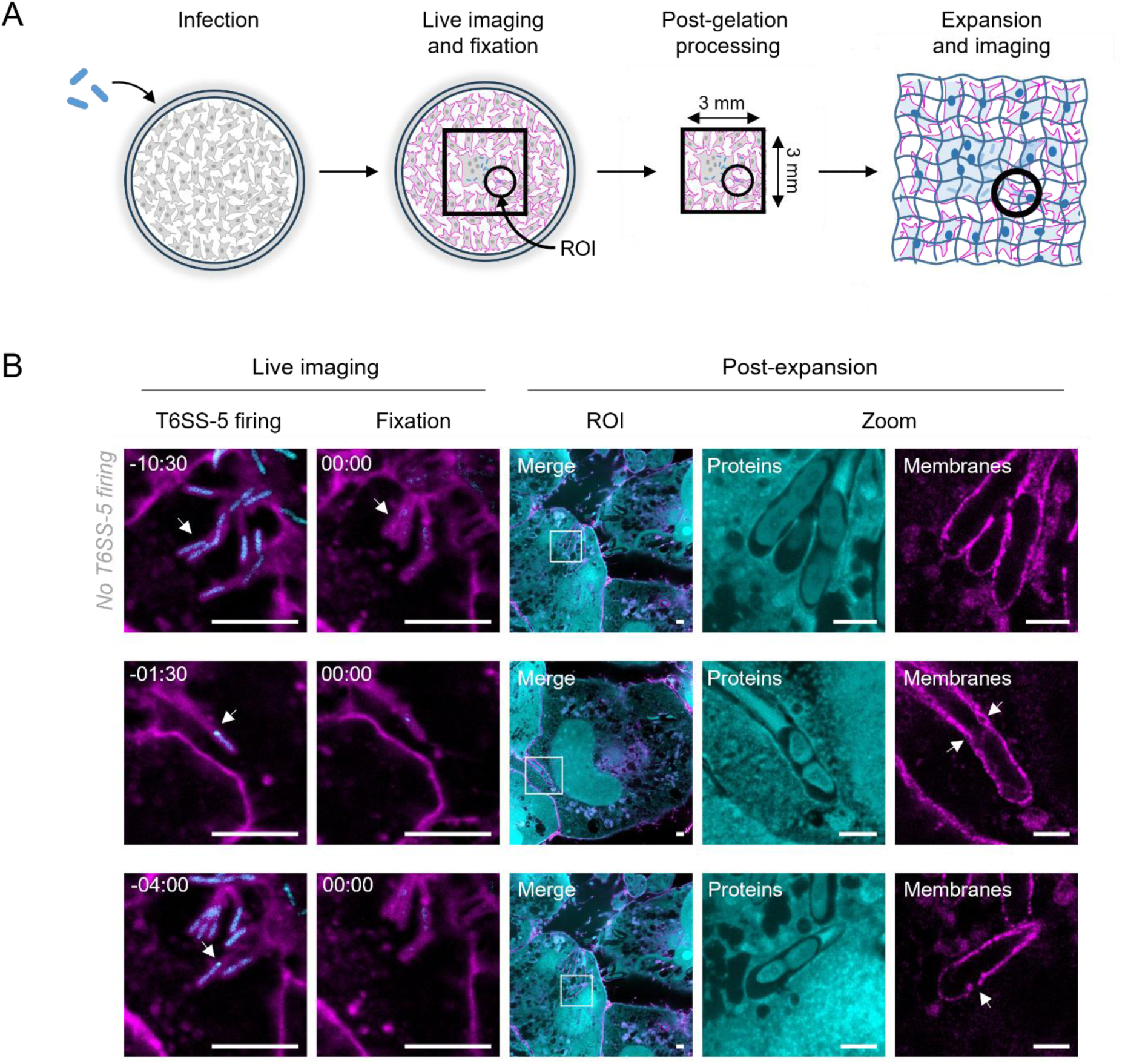
Use of LiFEx to correlate T6SS-5 assembly with membrane ultrastructure. **(A)** Schematic representation of the LiFEx pipeline. **(B)** Correlative live-cell and post-expansion imaging of bacteria-containing membrane protrusions. For example, live-cell images show the T6SS-5 firing event, when applicable (T6SS-5 firing), or the first frame of the timelapse, and the first frame acquired after fixation (Fixation). Scale bar, 10 μm. Post-expansion images show the corresponding region of interest (ROI), with the area indicated by the white square shown at higher magnification (Zoom). Examples include a bacterium with no detected T6SS-5 firing during the time-lapse acquisition (top), and bacteria that assembled T6SS-5 at 1min30sec (middle) or 4 min (bottom) before fixation. Top left numbers are time in min:sec. Scale bar, 10 μm (post-expansion).

At the end of the time-lapse acquisition, the cells were fixed directly on the microscope by introducing fixative from a syringe through a thin tube, thereby minimizing displacement of the stage or sample and preserving the position of the ROI. Following fixation, the entire marked area was scanned using a low-magnification objective (20×) to generate an overview image that could subsequently be used to generate a reference map for locating the cells after expansion. The sample was then removed from the microscope and processed using our improved expansion protocol as described above (Figure 2A), with one major modification: after complete gelation within the well, the gel was cut around the previously imaged 3×3 mm area, and only this small gel fragment was further processed for homogenization, staining, and expansion. This workflow allowed the region characterized by live-cell microscopy to be retained throughout sample processing and subsequently identified after expansion for high-resolution imaging.

Using the LiFEx pipeline, we first assessed how rapidly cellular dynamics were arrested following the addition of fixative. Importantly, movement of cellular and bacterial structures stopped within 30 s of addition of the fixation mixture, providing a defined temporal reference point for subsequent correlation with post-expansion images.

We then used LiFEx to follow bacteria-containing membrane protrusions for 10 to 30 min before fixation. Live-imaging allowed us to determine the time interval between T6SS-5 assembly and fixation, and subsequent post-expansion imaging showed the associated membrane organization (Figure 6B). We first examined protrusions containing bacteria in which no T6SS-5 activity was detected throughout the time-lapse acquisition. At high resolution, the labeling of the membranes surrounding these bacteria appear continuous (Figure 6B, first row). Interestingly, in a protrusion containing a bacterium that fired its T6SS-5 only 1.5 min before fixation, no clear membrane lysis was observed. However, small irregularities in the alignment of the two membranes forming the protrusion were visible (Figure 6B, second row). In contrast, when T6SS-5 firing occurred 4 min before fixation, the protrusion displayed a clear disruption of membrane integrity, characterized by an absence of the membrane labeling (Figure 6B, third row). Together, these observations suggest that membrane disruption may become apparent only minutes after T6SS-5 assembly.

## Discussion

Expansion microscopy is a powerful approach for super-resolution imaging of thick biological samples using confocal fluorescence microscopy. However, this approach can be challenging when applied to samples containing structurally distinct cell types, such as mammalian cells and bacteria. Here, we adapted the TREx protocol to expand mammalian cells infected with intracellular bacteria. This workflow enables approximately 9-fold expansion of the host-cell environment while preserving dense membrane labeling and pan-protein staining (Figure 2). Expansion enabled visualization of membrane organization associated with bacterial cell-to-cell dissemination and revealed membrane remodeling patterns that were not accessible using conventional fluorescence imaging (Figure 3).

A major challenge encountered during the development of our approach was the expansion difference between host cells and intracellular bacteria. Whereas host cells expanded approximately 9-fold, intracellular *B. thailandensis* expanded approximately 5.5-fold and showed heterogeneous expansion. We therefore investigated which bacterial features or experimental parameters could account for this incomplete and heterogeneous expansion. Unexpectedly, bacteria were still able to partially expand in the absence of PG degradation, indicating that the peptidoglycan layer does not completely prevent bacterial expansion (Figure 4). Nevertheless, expansion appeared more anisotropic under these conditions, consistent with the mechanical role of PG and with our observation that PG restricts proteinase K access to the bacterial cytoplasm. The influence of PG on bacterial expansion is consistent with previous ExM studies showing that PG hydrolysis increases expansion efficiency, but requires species-specific optimization^6^. However, in our experiments, PG degradation did not resolve the remaining expansion heterogeneity, and additional OM destabilization with EDTA did not substantially improve expansion (Figure S1). Heterogeneous expansion despite cell-wall degradation has also been reported for other bacterial species^6,17^, further indicating that the bacterial cell wall is not the only factor restricting complete and homogeneous expansion.

In contrast, fixation conditions had a pronounced effect on bacterial expansion patterns (Figure 5). The effect of fixation conditions on bacterial expansion has also been reported in a previous ExM study, in which the authors concluded that GA prevented efficient expansion of *Staphylococcus aureus*^7^. However, in our experiments, neither the absence of GA nor reducing the concentration of the fixatives prior to expansion improved bacterial expansion, but rather revealed a complex relationship between fixation conditions and bacterial expansion.

Importantly, the difficulty in achieving expected bacterial expansion with the TREx protocol is not restricted to our intracellular infection model. A recent application of the TREx protocol to planktonic *Escherichia coli* achieved only approximately 4-fold expansion in a single step and required iterative expansion to reach a final expansion factor of approximately 11-fold^19^. Thus, the limited expansion observed for intracellular *B. thailandensis* appears to reflect a broader challenge in applying high-expansion ExM protocols to bacteria, rather than a limitation specific to our approach or to intracellular bacteria.

Despite heterogeneous bacterial expansion, our approach enabled detailed visualization of the host membranes surrounding intracellular *B. thailandensis*. Dense membrane labeling revealed continuous and disrupted bacteria-containing protrusions as well as regular and locally distorted double-membrane structures. These morphologies may represent different states of membrane remodeling during bacterial dissemination, but ExM is a fixed-sample based technique and therefore cannot establish a temporal relationship between the patterns.

This limitation motivated the development of LiFEx, to address this gap by correlating live-cell events with the post-expansion architecture of the same cellular regions (Figure 6). Fixation directly during live imaging provided a defined endpoint, with detectable membrane dynamics and bacterial movement stopping within 30 s under our imaging conditions. Protrusions without detected T6SS-5 assembly displayed continuous membrane labeling after expansion, whereas increasing membrane alterations were observed in samples fixed 1.5 and 4 min after T6SS-5 activity was detected.

Taken together, LiFEx adds temporal context to the high-resolution structural information provided by ExM. This approach is not restricted to *B. thailandensis* or T6SS activity and could be applied to other dynamic cellular processes that can be identified by live-cell imaging and subsequently correlated with high-resolution cellular structures after expansion. This method will provide unprecedented insights into intracellular lifestyle of bacterial pathogens.

## Acknowledgments

We thank Alexia Loynton-Ferrand from the Imaging Core Facility (University of Basel) for support with data acquisition. The work was supported by Forschungsfonds of the University of Basel.

## Material and methods

### Bacterial strains and culturing conditions

*B. thailandensis* strain expressing TssB-5 fused to mScarlet-I was generated as previously described^26^. Frozen *B. thailandensis* stock was used to inoculate low-salt LB overnight at 37°C with aeration. The next day, this culture was diluted to 1:100 in fresh low-salt LB and grown until mid-log phase (OD_600nm_ = 0.3-0.4).

### Mammalian cell lines and culturing conditions

A549 lung cells (CCL-185, ATCC) were cultured in Ham’s F-12 media (Sigma-Aldrich) supplemented with 10% heat-inactivated fetal calf serum (FCS, BioConcept AG). All experiments were performed using cells from passages 2 to 15.

### Mammalian cell infection

Twenty-four hours prior to infection, A549 cells were seeded onto sterile 13-mm round coverslips, or µ-slide 8-well glass bottom (Ibidi GmbH, 80807) at 3.04 × 10^5^ or 8 × 10^4^ cells per well, respectively. Exponentially growing bacteria and mammalian cells were washed twice with OptiMEM Reduced Serum Medium with GlutaMAX supplement (Sigma-Aldrich) right before infection. *B. thailandensis* were added to the wells at a MOI of 50 or 100. The cells were then spun down for 5 min at 300 × g and placed at 37°C with 5% CO_2_. After 1 hour of incubation, infected cells were washed twice with OptiMEM supplemented with 300 μg/mL of kanamycin (OptiMEM-Kan) to kill extracellular *B. thailandensis*. The cells were further incubated in the presence of antibiotics for 17 hours. Infected cells were fixed with pre-warmed 4% PFA and 0.1% GA mix in PBS 1× for 10 min at 37°C, then rinsed 3 times with 1× PBS.

### Solution preparation for ExM

Sodium acrylate was either purchased (Sigma, 408220) or made by neutralizing acrylic acid as described previously^31^. Briefly, a 4 M sodium acrylate solution was generated by mixing 5.5 mL of acrylic acid and 4.5 mL of ddH2O. Next, 7.2 mL of 10 M NaOH was added gradually. The pH was then adjusted to 7.5-8 with 1 M NaOH. Finally, ddH_2_O was added to obtain a final volume of 20 mL. All the steps were performed under a fume hood in a beaker with a magnetic bar for constant agitation and on ice to prevent excessive heat during the reaction. The solution is then transferred to a Falcon tube and stored at -20°C.

The gelation solution was freshly prepared for each experiment. This solution contained 1.1 M sodium acrylate, 2 M acrylamide, 0.005% of N,N’-methylenebisacrylamide, PBS 1×, 0.15% of ammonium persulfate (APS) and 0.15% of *N*,*N*,*N*′,*N*′-Tetramethylethylenediamine (TEMED). TEMED and APS were added last to the monomer solution, right before use. All components were pre-cooled and mixed on ice.

For enzymatic homogenization of the sample, the gel was incubated in a solution containing 9 U/mL Proteinase K (Thermo Fisher, EO0491) in TAE buffer (40 mM Tris, 20 mM acetic acid, and 1 mM EDTA) supplemented with 0.5% Triton X-100 and 0.8 M guanidine-HCl.

mCLING-Atto647N (Synaptic Systems, 710 006AT1) and mCLING-Atto488 (Synaptic Systems, 710 006AT3) were resuspended into ddH_2_O by piercing the rubber cap with a syringe and vortexed thoroughly in all orientations to ensure complete resuspension of the powder before opening. The absorbance of the fluorophores was measured to estimate the quality of the batch, as high variability of membrane labeling quality was observed within batches.

### Optimized ExM protocol for intracellular bacteria study

ExM was applied to fixed infected mammalian cells as follows. The lipid membranes were labeled using 5 µM of mCLING-Atto647N (Synaptic Systems, 710 006AT1) or mCLING-Atto488 (Synaptic Systems, 710 006AT3) in PBS 1× overnight at room temperature. The following day, cells were permeabilized with 0.1% (v/v) Triton X-100 for 10 min at room temperature followed by three washes with PBS 1×. To degrade the PG, the sample was treated with 160 U/ml mutanolysin and 1 mg/ml lysozyme in 50 mM phosphate buffer supplemented with 1 mM MgCl2 for 3h at 37°C. The sample was then washed twice with PBS 1× and incubated with 100 µg/mL acryloyl-X SE (AcX) (Thermo Fisher, A20770) in PBS 1× overnight at room temperature. The next day cells were washed once with PBS 1× prior to gelation. For the gelation, silicone spacers (0.6 mm thick; Grace Bio-Labs Press-To-Seal silicone isolator, GBL665501) were customized to give the gels an irregular shape, which facilitated their orientation after expansion. The silicone spacers were attached to a parafilm layer at the bottom of a plastic box and placed on ice for 10 min to cool down. 100 µL of cold gelation solution (see recipes in the section “Solution preparation”) were placed on the parafilm and the coverslip was added on top, with the cells facing down to the gelation solution. After 5 min of incubation on ice, wet papers were added in the box before incubation for 1h at 37°C to allow gel polymerization. Afterwards, the sample was homogenized with 9 U/mL Proteinase K (Thermo Fisher, EO0491) in TAE buffer (containing 40 mM Tris, 20 mM acetic acid, and 1 mM EDTA) supplemented with 0.5% Triton X-100 and 0.8 M guanidine-HCl for 3h at 37°C with agitation. The gel was then washed three times with 1× PBS before incubation with 20 μg/mL fluorescent NHS ester ATTO 488, 594 or 647 (Sigma, 41698, 08741 and 07376) and 20 µM Hoechst 33342 (Thermo Fisher, 62249) in PBS 1× for 1 h at room temperature with agitation. Finally, the gels were washed 3 times with ddH_2_O and placed in a Petri dish filled with ddH_2_O. The ddH_2_O was exchanged twice after 30 min and the gels were left in water for expansion for at least 24 h at room temperature with agitation.

Note that when mentioned, cells were treated with 5 mM of EDTA in PBS 1× for 10 min after cell permeabilization with Triton.

### LiFEx

A549 cells were seeded in 1 cm² wells on a µ-slide 8-well glass bottom and infected with *B. thailandensis* at a MOI of 50, as described in the section “Mammalian cell infection” above. At 17 hpi, live-cell membranes were stained with 0.5 µg/mL of CellMask DeepRed (ThermoFisher, C10046) and 20 µM Hoechst 33342 (Thermo Fisher, 62249) in pre-warmed OptiMEM-Kan for 10 min at 37°C with 5% CO2 and washed 3 times with OptiMEM-Kan. The slide was then placed under a confocal spinning-disk microscope for live imaging at 37°C with 5% CO2. To facilitate further processing, a region of interest (ROI) was imaged live within a limited area of 0.09 cm² (3 mm × 3 mm), marked at the bottom of the well. A timelapse of the ROI was acquired with a 60× oil objective for 10 to 30 min with an interval of 30 s. The sample was then fixed directly under the microscope using a simple tubing system to preserve immobility of the ROI during imaging. A needle connected to a 1 mL graduated syringe with a 1 mm diameter tube (Masterflex® Transfer Tubing), the other end of which was placed in the well. The fixatives were directly injected into the cell culture medium (1:1 ratio) at a 2× concentration (8% PFA and 0.2% GA), for a final concentration of 4% PFA and 0.1% GA. After fixation was completed, an overview of the marked area was scanned with a 20× objective. The sample was then removed from the microscope, and expansion was performed as described above, with the exception that after gelation, the gel was cut to the 0.09 cm² (3 mm × 3 mm) area prior to homogenization. After expansion, the same cells were imaged.

### Live-imaging and gel imaging

Live-imaging was performed on a Nikon ECLIPSE Ti2 with the Crestoptics X-Light V3 spinning disk confocal system at 37°C with 5% CO2 incubation. Every 30 sec, a Z-stack of around 5 μm was acquired with a Z-step of 200 nm. The following objectives were used: 20× air (0.8 NA, 0.8 mm WD) and 60× oil (1.42 NA, 0.13 mm WD).

For expanded samples, the Nikon ECLIPSE Ti2 with the CrestOptics X-Light V3 spinning disk confocal system with a 60× silicone oil (1.3 NA, 0.3 mm WD) objective or the Olympus SpinSR (CSU-W1) spinning disk confocal system with 60× silicon objective (1.3 NA, 0.3 mm WD) were used. A Z-stack of about 20 µm was acquired with Z-steps of 200-500 nm.

## Supplementary figures

**Figure S1:**
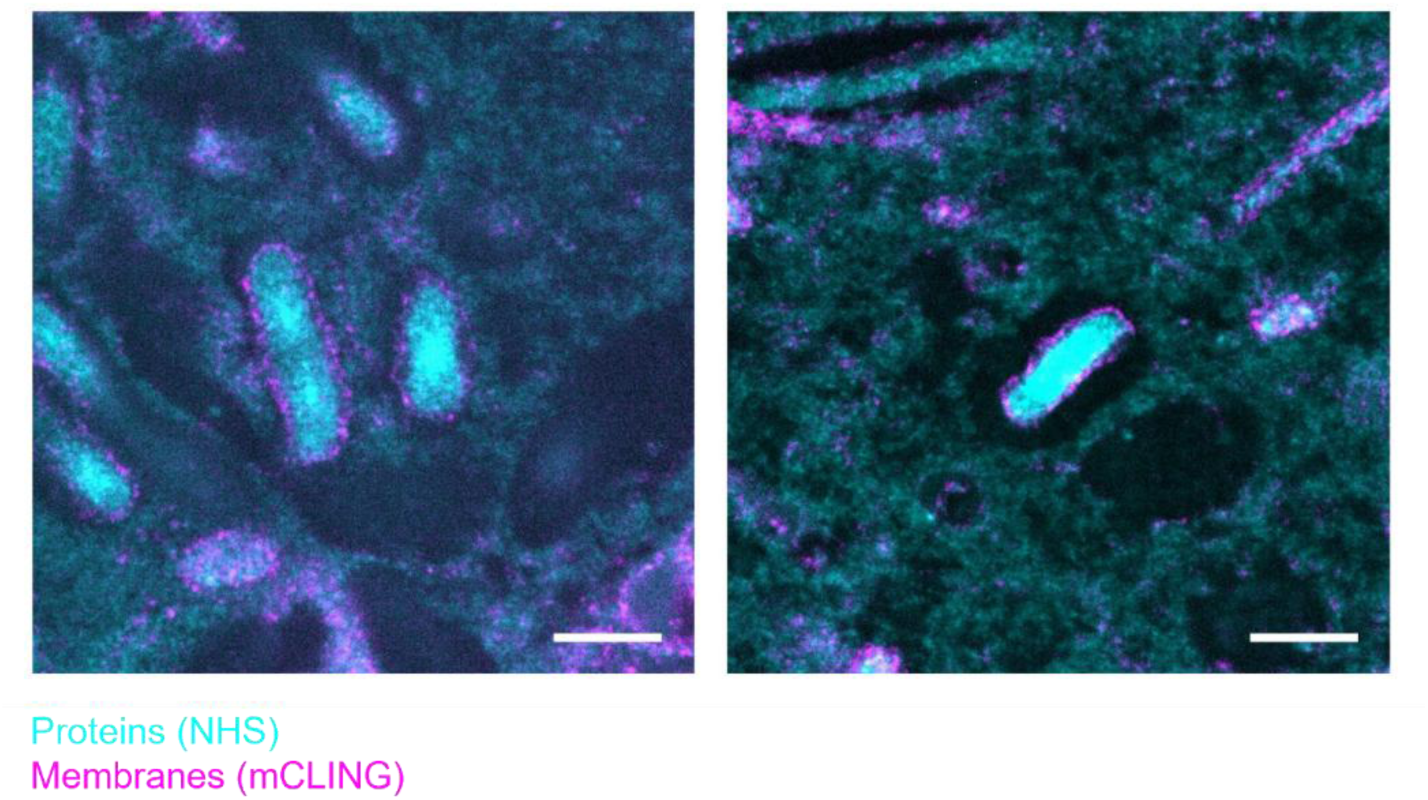
Heterogeneity of bacterial expansion. A549 cells are infected with *B. thailandensis* (MOI 100) for 17 h. After fixation, the ExM protocol is applied, and the expanded samples are imaged on spinning disk microscope. Membranes (mCLING) are shown in magenta, and proteins (NHS) in cyan. Scale bar, 10 µm (post-ExM).

**Figure S2:**
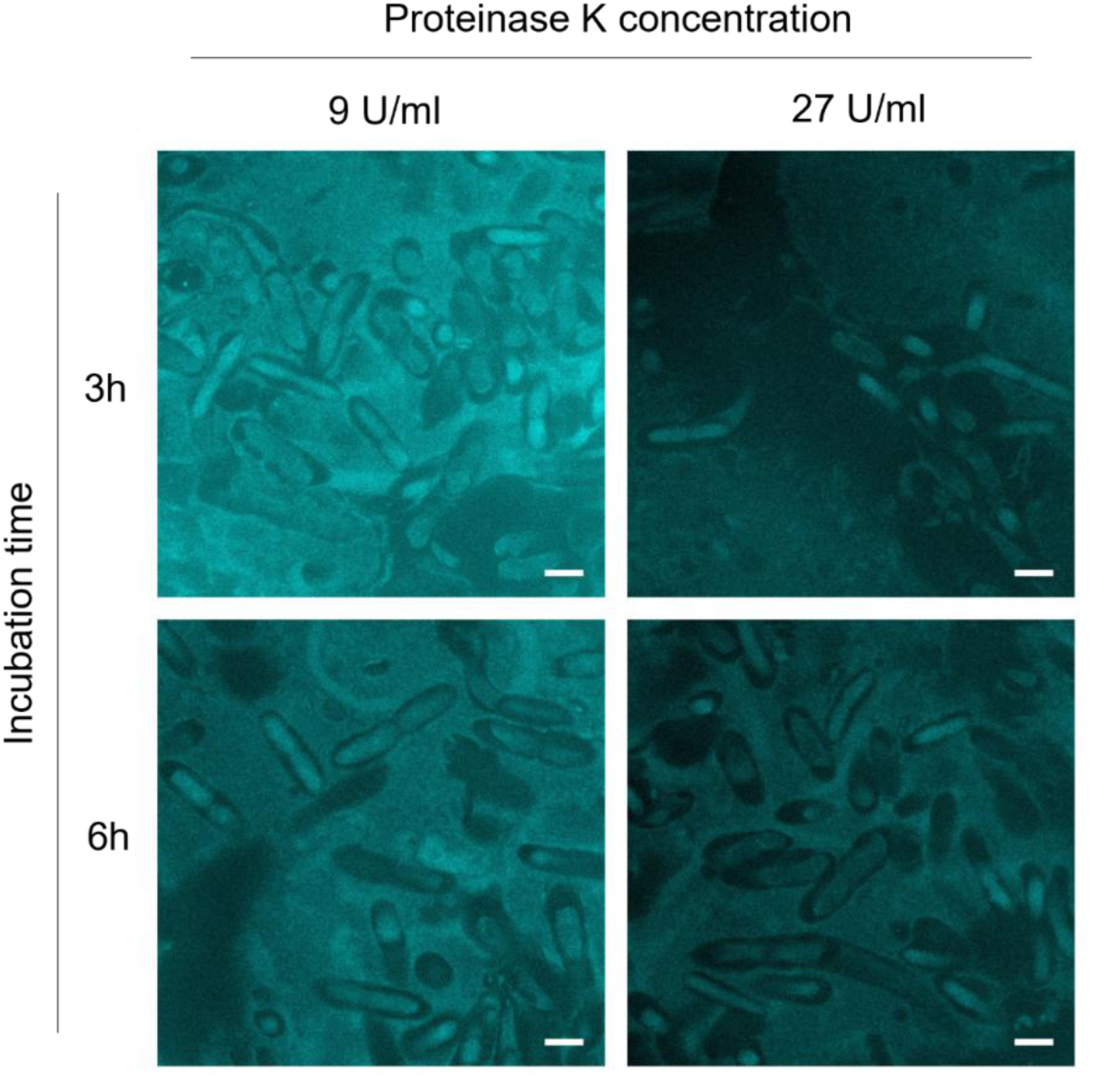
Effect of extensive proteolysis on bacterial expansion. A549 cells were infected with *B. thailandensis* (MOI 100) for 17 h. After fixation, the µTREx protocol was applied using 9 U/mL or 27 U/mL proteinase K for 3 h or 6 h. Protein staining (NHS) is shown in cyan. Scale bar, 10 µm (post-ExM).

